# CK1α, FAM83H, and FAM83B contribute to bundling of neurofilaments and are sequestered in cellular and mice models of ARSACS

**DOI:** 10.1101/2024.10.01.616079

**Authors:** Caitlin S. Atkinson, Zacharie Cheng-Boivin, Nancy Larochelle, Sandra Minotti, Benoit J. Gentil

**Affiliations:** Department of Kinesiology and Physical Education McGill University, Montreal, QC H3A 2B4, Canada; Department of Neurology and Neurosurgery and Montreal Neurological Institute, McGill University, Montreal, QC H3A 2B4, Canada

**Keywords:** ARSACS: Autosomal Recessive Spastic Ataxia of the Charlevoix Saguenay, IF: intermediate filaments, NF: neurofilaments, NFH: neurofilament heavy polypeptide, NFM: neurofilament medium polypeptide, NFL: neurofilament light polypeptide, ULFs: unit-length filaments, CK1: Casein Kinase 1, FAM83: FAMily with sequence similarity 83, DUF: Domain of Unknown Function

## Abstract

Autosomal recessive spastic ataxia of the Charlevoix-Saguenay (ARSACS) is a rare neurodegenerative disorder characterized by mutations in the *SACS* gene that encodes for the sacsin protein. Sacsin dysfunction in ARSACS results in neurofilament bundling, a phenotype observed in various cellular models of ARSACS. The mechanisms underlying bundling in ARSACS remain unclear. With neurofilament phosphorylation controlling several processes of intermediate filament dynamics, its dysregulation may play a role in ARSACS. Accordingly, we investigated the interaction between CK1α (Casein Kinase 1α) and its adaptor proteins FAM83H or FAM83B (FAMily with sequence similarity 83). Here, we report that these target proteins are upregulated in ARSACS patient fibroblasts and co-localize at the sites of intermediate filament bundling. In the *Sacs*^*-/-*^ cerebellum, FAM83B compensated for the lack of FAM83H expression, suggesting a cell-type specific activity of CK1α that depends on the relative expression of its adaptor proteins. Further, CK1α inhibition with D4475 or knockdown of the target proteins with the CRISPR system caused neurofilament bundling, a phenotype that was only partially remedied by CK1α activation with SSTC3. Our findings suggest that CK1α, FAM83H, and FAM83B contribute to neurofilament bundling in ARSACS; however, the inability of CK1α to resolve neurofilament bundling may reflect an error in a priming phosphorylation event in ARSACS. Future research is needed to understand the hierarchical phosphorylation cascade to CK1α activity and its contribution to ARSACS pathology.

**Highlights:**

- CK1α and its cell-specific adaptors are upregulated in ARSACS and targeted to the sites of bundles
- Knockdown of CK1α, FAM83H, and FAM83B cause neurofilament bundling
- Activation of CK1α partially remediates bundling

## Introduction

Autosomal recessive spastic ataxia of the Charlevoix-Saguenay (ARSACS) is a progressive neurodegenerative disease characterized by a common group of symptoms that include cerebellar ataxia, spasticity, and peripheral neuropathy (1). ARSACS is caused by mutations in the *SACS* gene that lead to the loss of function of sacsin, a multi-domain protein enriched in neurons (2,3). Sacsin’s functions are unclear; however, studies into the functions of sacsin’s domains, along with omics approaches in ARSACS models, demonstrate a role in protein chaperoning and homeostasis (1,2). One of the first and most striking phenotypes described in cellular models of ARSACS is the bundling of neurofilaments (NFs), the intermediate filaments (IF) of the neuronal cytoskeleton (4). These NF bundles are accompanied by the hypophosphorylation of NF proteins in neurons of sacsin-deficient mice (*Sacs*^*-/-*^ *and Sacs*^*272/272*^) (4–6). Several fundamental cellular processes are disrupted because of these bundles (7), suggesting that the altered cytoskeletal organization of NFs may contribute to the development of the associated neurological symptoms. This bundling is replicated in many different cellular models of ARSACS (fibroblasts, neurons, SW13^*vim-*^, and astrocytes) carrying different types of IFs (4).

IFs are cell-specific elements of the cytoskeleton and consist of keratins, vimentins, and NFs. The assembly of IF networks follow similar stepwise processes that are well documented (8). NFs are heteropolymers composed of three main monomers: NFL, NFM, and NFH (9). These monomers have a similar N-terminus and α-helical core but differ extensively by their C-termini (9). Of these monomers, NFL is a core protein necessary for the formation of filamentous structures. These proteins associate into a dimer, which in turn polymerizes with another dimer to form an anti-parallel staggered tetramer through the association of their α-helical core domain (10). The resulting tetramer polymerizes with another similar tetramer to form a fundamental unit or “unit-length filaments” (ULFs) of the mature filament, which can be subsequently added or removed (10,11). The ULFs anneal end-to-end until they form the mature IF that can be many microns in size (11,12).

While the mechanisms underlying the compaction or straightening processes of NFs remain unclear, post-translational modifications such as phosphorylation play a role in IF assembly and transport within the cell (13,14). Phosphorylation is primarily controlled by kinase proteins, such as the Casein Kinase 1 (CK1) family. Notably, a multi-omic analysis identified these proteins to have upregulated expression in *Sacs*^*-/-*^ neuroblastoma cells (15). However, the role of CK1 in the ARSACS molecular pathogenic cascade is unknown. CK1 is a family of seven constitutively active isoforms that contain similar and conserved N-termini while their C-termini differ (16,17). This family of proteins has a wide variety of functions, including cytoskeletal integrity, circadian rhythm, and cellular differentiation (18). Each isoform has a kinase domain that phosphorylates multiple sites but primarily phosphorylates serine/threonine residues at the *n*-3/4 position (19). Inhibition of the specific isoform, CK1α, was shown to be responsible for NF bundling and hypophosphorylation of NFH in NB2a/d1 neuroblastoma cells (15).

How these CK1 isoforms specifically recognize and associate with their targets was previously unknown, until the recent identification of FAMily with sequence similarity 83 (FAM83) proteins. FAM83 is a family of eight proteins that are named accordingly from A to H (18). The functions of this family of proteins are poorly understood; however, all eight FAM83 members have the same domain at the N-terminus known as DUF1669 (Domain of Unknown Function 1669), which was recently characterized to be a major anchor of the CK1 α, δ, and ε isoforms with their respective FAM83 proteins (16,17). Aside from their DUF1669 domain, all members of this family have distinct amino acid sequences at their C-termini, which may confer specific binding to different partners (16).

Among the eight proteins, FAM83H and FAM83B contain the most conserved sequences at their C-termini, and have been shown to interact with CK1α isoforms (17). In particular, FAM83H recruits CK1α to regulate keratin filament networks in colorectal cancer and ameloblastoma cells (18,19). Indeed, knockdown of FAM83H induces keratin filament bundling while overexpression promotes keratin filament disassembly (19). Similarly, the inhibition of CK1α induces keratin filament bundling and reverses keratin filament disassembly, which suggests that both proteins are dependent upon each other (18). On the other hand, the role of the interaction between FAM83B and CK1α on IF dynamics remains poorly characterized. However, FAM83B has been associated with modulating the vimentin network in cancer cells, whereby *FAM83B* knockdown increased mesenchymal vimentin expression to promote cell migration (20). Although most research to date has focused on the role of FAM83B in oncogenic signaling, the conserved C-termini between FAM83H and FAM83B may suggest a common role in cytoskeletal organization. In turn, their interaction with CK1α may contribute to the altered NF network in ARSACS. With the described interaction between FAM83 and CK1 protein families, we aimed to investigate the relationship between CK1α and the two adaptor proteins FAM83H and FAM83B and their effect on the formation and organization of NFs in various models of ARSACS mutations.

## Methods

### Animals and ethic statement

*Sacs*^*-/-*^ and wild-type (*Sacs*^*+/+*^) C57BL mice as previously described (21) were bred in our animal facility. Experiments were approved by the Animal Care Committee of the Montreal Neurological Institute (McGill University protocol #7437) and all procedures followed the Canadian Council on Animal Care guidelines. Secondary use of human-derived cells was approved by McGill ethics committee (A07-M34-19B).

### Cell Culture

Human dermal fibroblasts were obtained from the CellBank Repository for Mutant Human Cell Strains (McGill University Health Complex, Montreal, QC, Canada). Immortalized fibroblasts from control (MCH74) and an ARSACS patient expressing the homozygous ARSACS-causing p.2801delQ variant (7373) previously described in (4,22), were cultured in DMEM with 10% fetal bovine serum. To modulate CK1α activity, cultures were treated with 100 μM D4476 (HY-10324, MedChemExpress) in DMSO (0h, 3h, 24h) or 1 μM SSTC3 (HY-120675, MedChemExpress) in DMSO (0h, 6h, 24h) prior to immunolabelling.

SW13^vim-^ cells, which lack endogenous IFs, were cultured in Dulbecco’s Modified Essential Media (DMEM) with 10% fetal bovine serum (15). To evaluate NF assembly, SW13^vim-^ cells were co-transfected with plasmids encoding wild-type *NEFL* and *NEFH*, along with CRISPR-Cas9 knockout plasmids encoding control guide RNA or guide RNA targeting *FAM83H* (sc-415452, Santa Cruz Biotechnology), *FAM83B* (sc-415158, Santa-Cruz Biotechnology) or *CK1α* (sc-400898-KO-2, Santa Cruz Biotechnology) and EGFP, using Lipofectamine 2000 as previously described in (23). 48 hours after transfection, indirect immunofluorescence detection of NFL was used to evaluate the NF network in transfected cells, as identified by EGFP expression.

Primary cultures of dissociated spinal cord-dorsal root ganglia were prepared from C57Bl6 E13 *Sacs*^-/-^ and *Sacs*^*+/+*^ mice and were maintained in Eagle’s Minimal Essential Media (EMEM) as previously described (24). Cultures were used six weeks following plating to allow neuronal maturation and NF bundle formation in *Sacs*^-/-^ motor neurons. Cultures were treated with D4476 as described in the fibroblasts. Because motor neurons in long-term primary spinal cord cultures are not transfectable, CRISPR-Cas9 control and *CK1α* knockout plasmids were expressed by intranuclear microinjection in *Sacs*^*+/+*^ motor neurons in culture as described in (25). 48 hours after microinjection, indirect immunofluorescence detection of NFL was used to assess the NF network in microinjected cells, as identified by EGFP expression.

### Immunohistochemistry analysis

Two female and two male adult mice (>200 days old) per group were anaesthetized by isoflurane and euthanized by decapitation. The whole brains were removed and transferred to 10% formalin for 4 days. The fixed brains were processed, paraffin embedded, and sectioned at the Rosalind & Morris Goodman Cancer Institute at McGill University. For immunohistochemistry analyses, sagittal sections were deparaffinized with Xylene then underwent a sodium citrate heated antigen retrieval process. The slides were blocked in 10% Horse Serum, 3% BSA, and 0.3% Triton-X in TBS for 1-2 hours at room temperature, then incubated with primary antibodies diluted in blocking buffer overnight at 4° C. The primary antibodies included anti-FAM83H (HPA024505; Sigma-Aldrich), anti-FAM83B (ab122175; abcam), anti-CK1α (ab206652; abcam), and anti-NFL (clone NR4, N5139, Sigma-Aldrich). The following day, slides were washed three times with TBS and incubated with the appropriate Cy2 and Cy3 conjugated secondary antibodies (Jackson ImmunoResearch) along with DAPI for 1-2 hours at room temperature. The slides were washed and mounted with coverslips using Immunomount and left to dry overnight. Imaging was performed on a Zeiss Observer.Z1 microscope with the ZenBlue 2.6 Software (Carl Zeiss Canada Ltd.).

### Immunofluorescence

Indirect immunofluorescence was carried out after fixation with 4% Paraformaldehyde, using anti-vimentin (clone V9, MA5-11883, Thermo Fisher), anti-NFL (clone NR4, N5139, Sigma-Aldrich), anti-FAM83H (HPA024505; Sigma-Aldrich), anti-FAM83B (ab122175; abcam), and anti-CK1α (ab206652; abcam) primary antibodies. Corresponding Cy2 and Cy3 conjugated secondary antibodies were used (Jackson ImmunoResearch) along with DAPI. Images were captured using a Zeiss Observer.Z1 microscope with the ZenBlue 2.6 Software (Carl Zeiss Canada Ltd.).

### Data analysis and statistics

Images were analyzed using Fiji software (26). To evaluate the expression levels of FAM83H, FAM83B, and CK1α, the mean background adjusted fluorescence of each cell was determined for >50 cells per condition from >2 independent replicates. For analyses of vimentin and NFL networks, the percentage of cells carrying a normal (i.e., filamentous) versus bundled IF network was quantified for >3 independent replicates with >50 cells per replicate. Data are presented as means ± S.D. Statistical analysis was conducted using GraphPad Prism software, via two-tailed student’s t-test or one-way ANOVA with Tukey’s HSD as a posthoc test. Statistical significance was established at p<0.05.

## Results

### CK1α and FAM83B co-localize with NFL at the sites of bundles in the Sacs^-/-^ cerebellum

To determine which FAM83 proteins associate with NF proteins in the cerebellum, we investigated the subcellular distribution of CK1α, FAM83H, and FAM83B in relation to NFL using indirect immunofluorescence in >200-day-old (symptomatic) *Sacs*^*-/-*^ and *Sacs*^*+/+*^ mice. This brain region is severely affected in ARSACS, as characterized by Purkinje neuron degeneration that leads to cerebellar ataxia (27,28), with a prominent presence of NF bundles in these neurons. In the *Sacs*^-/-^ cerebellum, FAM83B was present in the granular layer, Purkinje cell layer, and the molecular layer, where it strongly co-localized with NFL bundles (Figure 1A). Conversely, FAM83H was barely detectable in Purkinje neurons (Supplemental Data 1) suggesting a cell-type specific expression of these two FAM83 proteins. CK1α was particularly abundant in the soma of both *Sacs*^*+/+*^ and *Sacs*^*-/-*^ Purkinje neurons *in vivo* (Figure 1B), while NFL was distributed in the axons of the granular layer in *Sacs*^*+/+*^ cerebellum mice and co-localized with CK1α in the cell body and the dendritic arborization of *Sacs*^*-/*-^ Purkinje neurons. (Figure 1). Overall, the co-localization of NFL bundles with FAM83B and CK1α in *Sacs*^-/-^ mice suggest that these proteins are targeted to the NFL bundling sites, yet they cannot execute their functions to remediate bundling.

**Figure 1.**
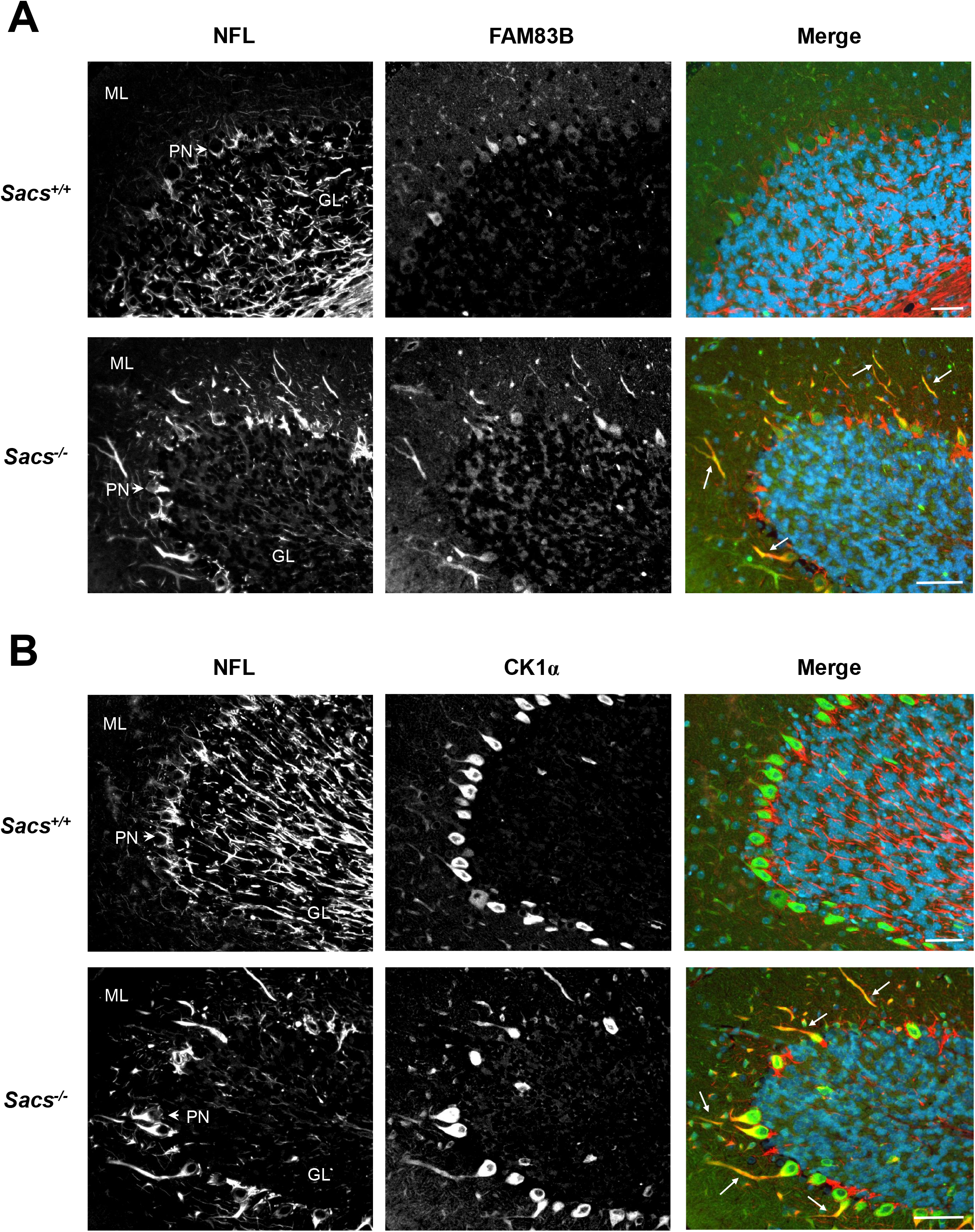
Co-localization of NFL with FAM83B and CK1α in the adult mouse cerebellum. The co-localization NFL and FAM83B or CK1α was assessed via immunohistochemistry in *Sacs*^*+/+*^ and *Sacs*^-/-^ cerebellar tissue. **A**. Representative images of double immunolabelling of NFL and FAM83B in the *Sacs*^*+/+*^ and *Sacs*^-/-^ cerebellum. In both genotypes, FAM83B is primarily expressed in the molecular layer of the cerebellum; in *Sacs*^*-/-*^ mice, FAM83B has greater expression in the granular layer and strongly co-localizes with the neurofilament bundles (white arrows). *n* = 4 mice per condition. **B**. Representative images of double immunolabelling of NFL and CK1α in the *Sacs*^*+/+*^ and *Sacs*^-/-^ cerebellum. In both genotypes, CK1α is highly expressed in Purkinje neurons. In *Sacs*^*-/-*^ mice, CK1α has greater co-localization with the neurofilament bundles (white arrows). *n* = 4 mice per condition. ML = molecular layer; PN = Purkinje neurons; GL = granular layer. Scale bar = 20 µm.

### CK1α, FAM83H, and FAM83B co-localize with vimentin bundles in fibroblasts derived from ARSACS patients

To determine whether the mislocalization of FAM83B and CK1α with IF bundles is specific to ARSACS, we investigated the subcellular distribution of FAM83H, FAM83B, and CK1α in fibroblasts derived from an ARSACS patient (Line 7373). As shown in Figure 2, the vimentin network was bundled in 7373 fibroblasts while MCH74 control fibroblasts showed a normal vimentin network as previously described (29). FAM83H, FAM83B, and CK1α co-localized with vimentin, which appeared to be increased in 7373 fibroblasts and concentrated at the sites of vimentin bundling (Figure 2A-C). In contrast to our findings in Purkinje neurons, FAM83H strongly aligned with these vimentin bundles (Figure 2A).

**Figure 2.**
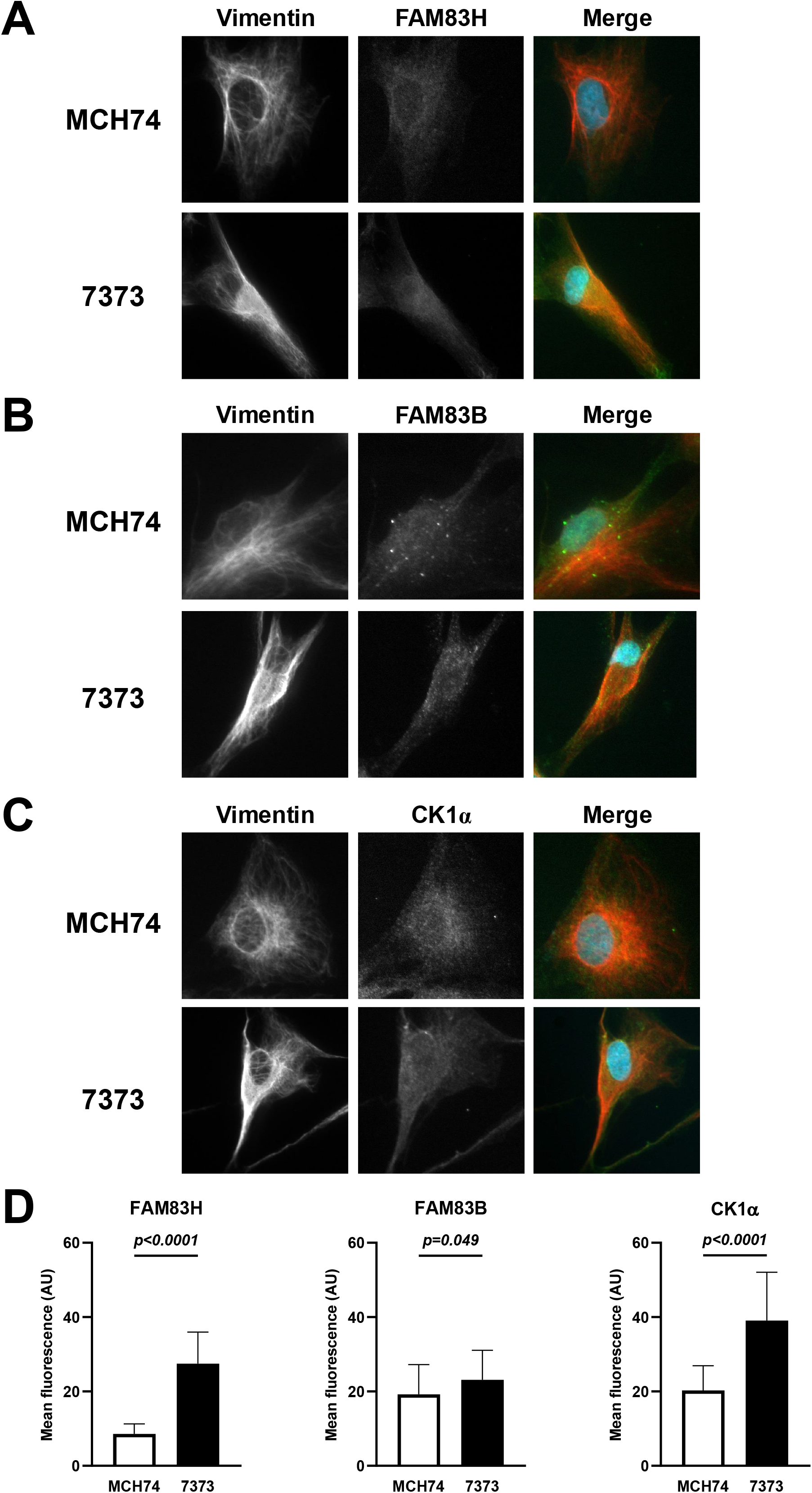
Co-localization of vimentin with FAM83H, FAM83B and CK1α in healthy and ARSACS human fibroblasts. The co-localization of vimentin with FAM83H, FAMB83B, and CK1α was assessed via indirect immunofluorescence in human dermal fibroblast from control (MCH74) and ARSACS (7373) donors. **A**. Representative images of double immunolabelling of vimentin and FAM83H **B**. vimentin and FAM83B, **C**. vimentin and CK1α. **D**. Quantification of the fluorescence intensity of FAM83H, FAM83B, and CK1α in MCH74 in comparison to 7373 fibroblasts. All proteins were significantly upregulated in 7373 fibroblasts. Background adjusted mean fluorescence intensity per cell is presented as condition means ± S.D (*n* > 50 cells/condition), with the *p*-values determined by two-tailed unpaired student’s *t*-test as labelled.

Quantification of the total fluorescence intensities of FAM83H, FAM83B, and CK1α in MCH74 and 7373 fibroblasts indicated an upregulation of protein levels in 7373 fibroblasts (Figure 2D). Specifically, FAM83H and CK1α were respectively upregulated 3-fold and 2-fold relative to MCH74, while FAM83B was upregulated by 20% (Figure 2D). Together, these results indicate that sacsin influences the localization and levels of CK1α, FAM83H, and FAM83B.

### CK1α, FAM83H, and FAM83B knockout causes bundling during NFL assembly in SW13^vim-^ cells

Next, we aimed to determine the contribution of FAM83H, FAM83B and CK1α in bundling during IF assembly. Adrenocarcinoma SW13^vim-^ cells lack IF proteins, making them a good model to study the *de novo* assembly of NFL (23). Therefore, we co-transfected plasmids encoding NFL, along with CRISPR-cas9-EGFP knockout systems targeting *FAM83H, FAM83B*, or *CK1α* in SW13^vim-^ cells, and assessed the NFL network morphology in EGFP positive cells. In SW13^vim-^ cells transfected with a plasmid carrying a control guide RNA, 23.6±1.1% of the cells demonstrated the bundling phenotype in their NFL network organization after 48 hours (Figure 3B). However, the percentage of cells presenting NFL bundles increased to 70.7±4.3% with *FAM83H*, 70.0±2.2% with *FAM83B* and 80.3±4.6% with *CK1α* knockouts (Figure 3B). Qualitatively, NFL bundles produced by *FAM83H, FAM83B* and *CK1α* knockouts were more packed than the basal bundling observed when using control guide RNA (Figure 3A). In addition, the NFL bundles were displaced to the periphery of the cell’s nucleus (Figure 3A). *CK1α* knockout increased the percentage of SW13^vim-^ cells carrying NFL bundles to the greatest extent, suggesting that CK1α is essential for *de novo* NFL formation (Figure 3B).

**Figure 3.**
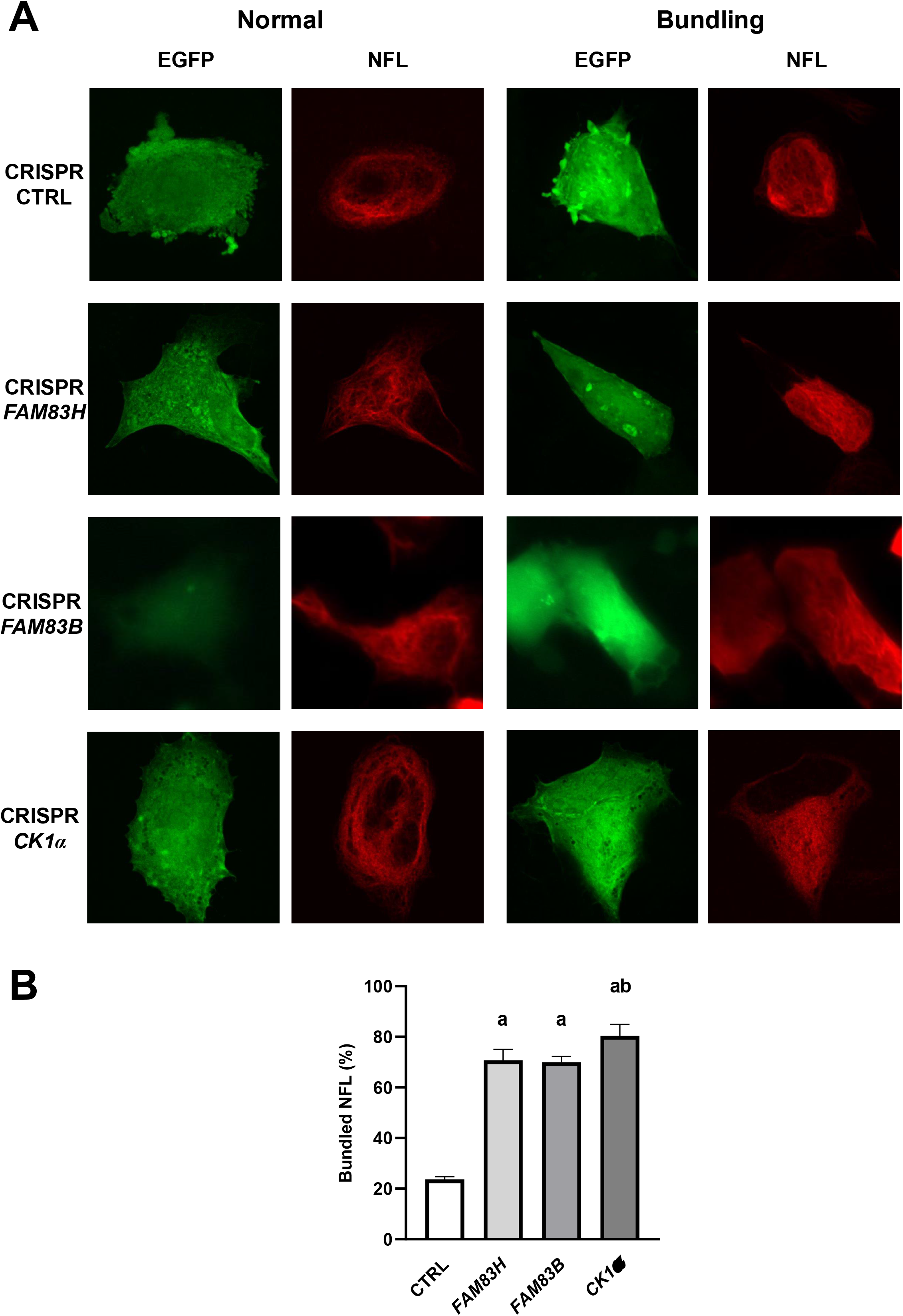
The effect of FAM83H, FAM83B, and CK1α knockout in SW13^vim-^ cells on neurofilament assembly. SW13^vim-^ cells were co-transfected with a plasmid carrying the CRISPR-Cas9 knockout plasmid targeting FAM83H, FAM83B, or CK1α and expressing EGFP, along with plasmids encoding NFL and NFH. **A**. Images show immunolabelling of the assembled NFL network. In the “NORMAL” column, images are representative of a normal NFL network in the different experimental conditions. In the “BUNDLING” column, images are representative of abnormal NFL network assembly, with a characteristic bundling of filaments. **B:** Quantification of the effect of gene knockout on neurofilament assembly. Shown is the percentage of total cells presenting with a bundled neurofilament network in the different experimental conditions (i.e. CRISPR-CTL, CRISPR-*FAM83H*, CRISPR-*FAM83B*, and CRISPR-*CK1α*). *FAM83H, FAMB83B*, and *CK1α* gene knockout significantly increased the proportion cells presenting a bundled NFL/NFH network. Data are expressed as means ± S.D (*n*=5 replicates/condition), with significant differences labelled a = *p*<0.05 versus control, b = *p*<0.05 versus FAM83H/FAM83B, as determined by one-way ANOVA with Tukey’s HSD post-hoc analysis.

### Treatment with the specific CK1 inhibitor, D4476, causes bundling in a time-dependent manner

To further investigate the role of CK1α in the formation of IF bundling, we pharmacologically inhibited CK1α activity with 100μM D4476 in cellular models of ARSACS and evaluated the IF network by indirect immunofluorescence at 3h and 24h post-treatment. First, treatment with D4476 caused an increase in the percentage of MCH74 fibroblasts with vimentin IF bundles in a time dependent manner (Figure 4). Qualitatively, these bundles are not different from the one observed in ARSACS 7373 fibroblasts (Figure 4A). D4476 treatment exacerbated the ARSACS phenotype by increasing the percentage of vimentin IF bundling in 7373 fibroblasts (Figure 4B). However, this percentage reached a maximum IF bundling (60.8±1.0%) at 3h post-treatment, with no further increase observed at 24 hours (Figure 4B).

**Figure 4.**
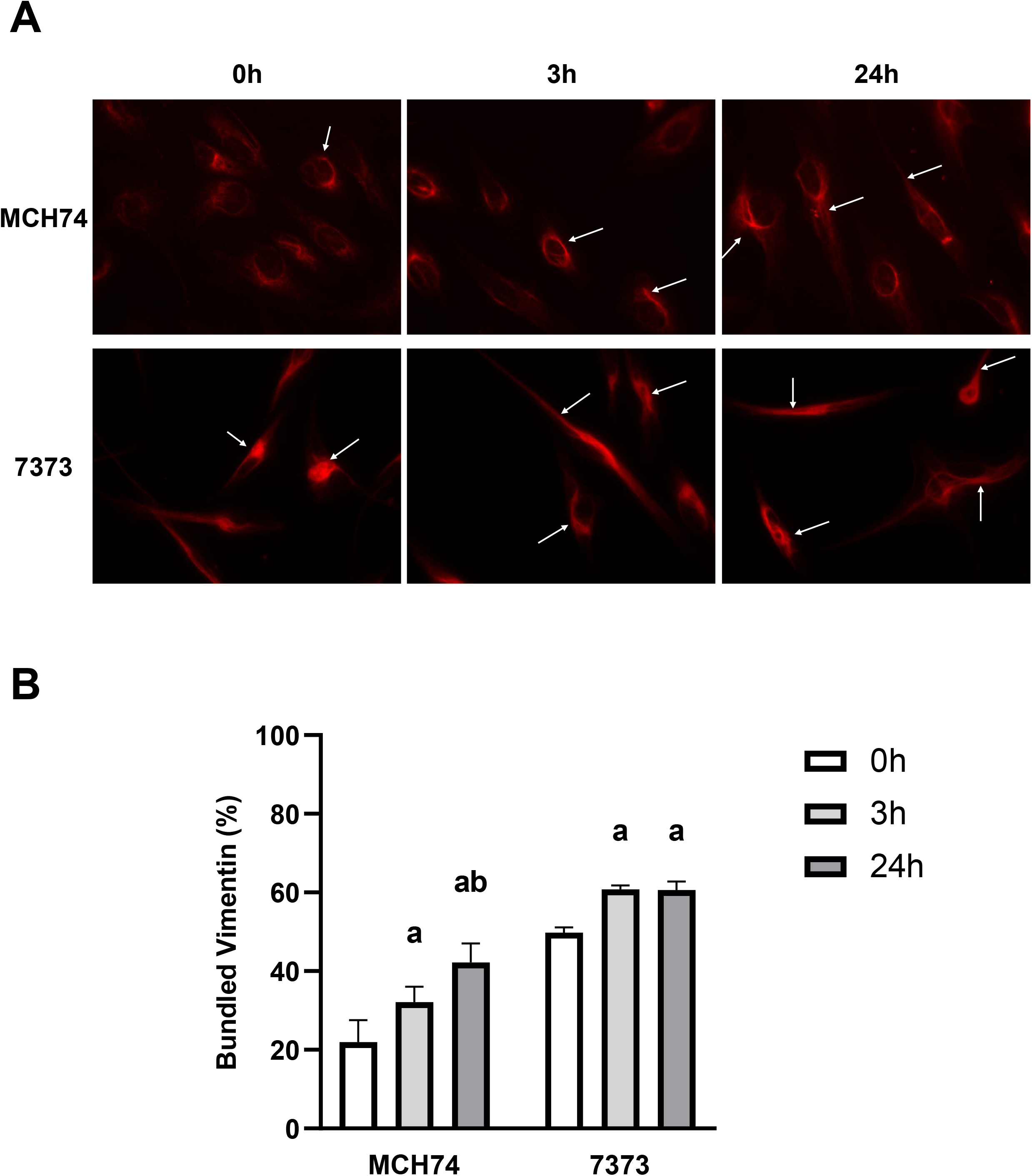
The effect of the CK1α inhibitor D4476 on vimentin bundling in healthy and ARSACS human fibroblasts. Human dermal fibroblast from control (MCH74) and ARSACS (7373) donors were treated with with 100 μM D4476, a CK1 inhibitor, for 0h, 3h, or 24h and the vimentin network was assessed via indirect immunofluorescence. **A**. Shown are representative images of the vimentin network of fibroblasts, with bundles highlighted by arrows. **B**. Quantification of the effect of D4476 on vimentin bundling in fibroblasts. Shown are the percentage of cells presenting with normal vimentin network in the different experimental conditions. D4476 significantly increased the proportion cells presenting a bundled vimentin network. Data are expressed as means ± S.D (*n*=3 replicates/condition) with significant differences between time-points within cell types labelled a = *p*<0.05 versus 0h and b = *p*<0.05 versus 3h, as determined by one-way ANOVA with Tukey’s HSD post-hoc analysis.

Next, we tested the effect of CK1α inhibition on pre-existing NF bundles in cultured *Sacs*^*-/-*^ murine motor neurons, as these neurons form well-established bundles in long-term cultures (30). *Sacs*^*+/+*^ motor neurons in culture expressed CK1α (Supplemental Data 2). D4476 treatment (100μM) caused NFL bundling in six-week-old *Sacs*^*+/+*^ motor neurons in a time-dependent manner, as shown by the increase in the percentage of motor neurons carrying NFL bundles (Figure 5). Qualitatively, these NFL bundles are not different from pre-existing NFL bundles observed in six-weeks-old *Sacs*^*-/-*^ motor neurons (Figure 5A). Knockout of *CK1α* expression in six-week-old *Sacs*^+/+^ murine motor neurons without pre-existing NFL bundles in culture led to the formation of NFL bundles (Supplemental Data 3), indicating that both lack of CK1α and inhibition of its enzymatic activity have a similar effect on the formation of NFL bundles. Contrary to 7373 fibroblasts, NFL bundling was exacerbated following D4476 treatment and did not reach a plateau after three hours in six-week-old *Sacs*^*-/-*^ motor neurons in culture (90% in motor neurons versus 60% in 7373 fibroblasts at 24 hours post-treatment), which suggests that a compensatory mechanism may protect 7373 fibroblasts, increasing the susceptibility of neurons.

**Figure 5.**
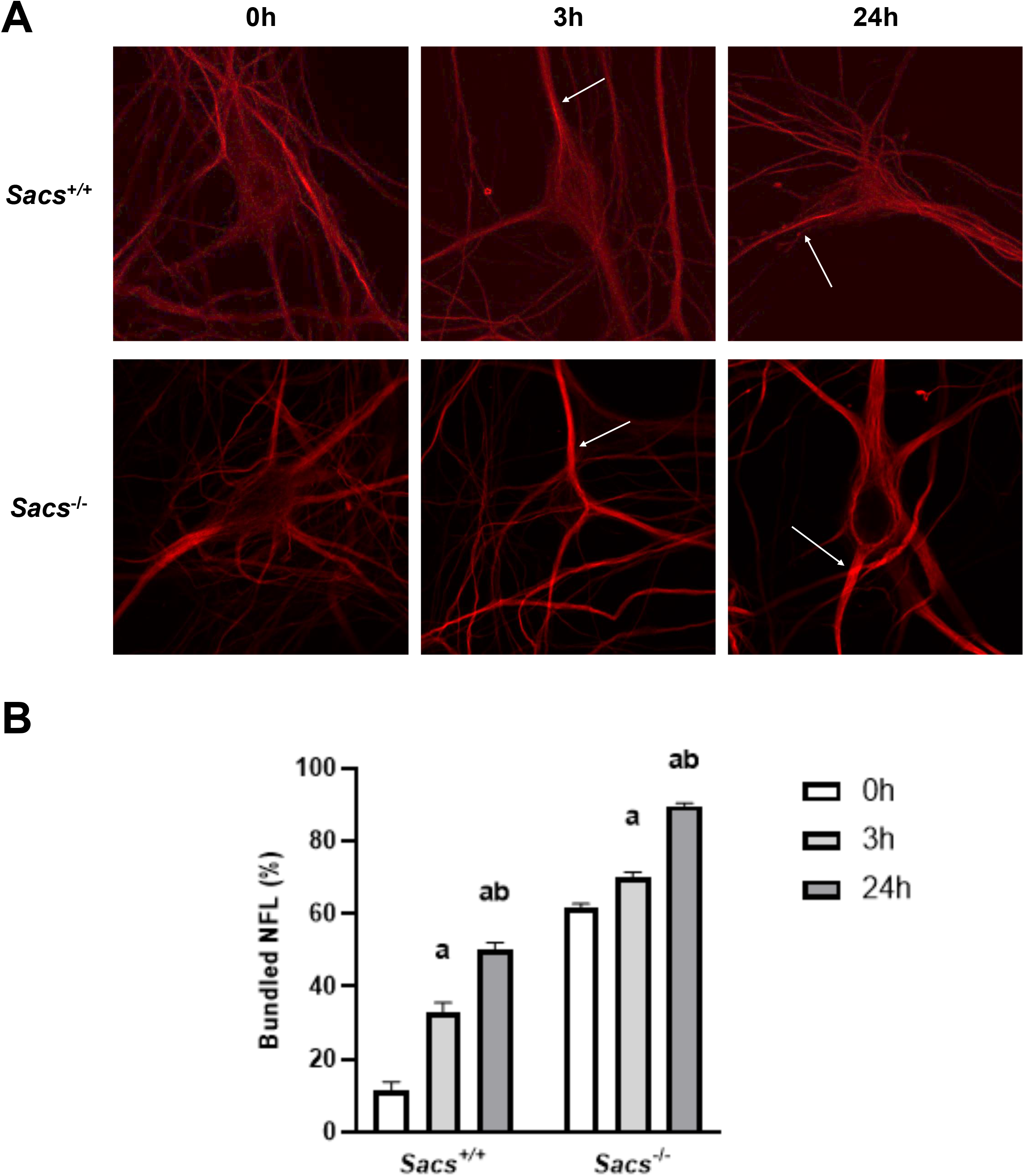
The effect of the CK1α inhibitor D4476 on neurofilament bundling in cultured *Sacs*^*+/+*^ and *Sacs*^-/-^ mouse motor neurons. Spinal cord-dorsal root ganglia primary cultures from *Sacs*^*+/+*^ and *Sacs*^*-/-*^ mice were treated with 100μM D4476, a CK1α inhibitor, for 0h, 3h, or 24h and the NFL network assessed via indirect immunofluorescence. **A**. Shown are representative images of the NFL network of motor neurons in different conditions, with bundles highlighted by arrows. **B**. Quantification of the effect of D4476 on NFL assembly in motor neurons. Shown is the percentage of cells presenting a normal NFL network in the different experimental conditions. D4476 significantly increased the proportion cells presenting a bundled NFL network. Data are expressed as means ± S.D (*n*=3 replicates/condition) with significant differences between timepoints within cell types labelled a = *p*<0.05 versus 0h and b = *p*<0.05 versus 3h, as determined by one-way ANOVA with Tukey’s HSD post-hoc analysis.

### A CK1α activator, SSTC3, minimally resolves neurofilament bundles

To investigate if the activation of CK1α can effectively resolve the abnormal vimentin IF bundling, we treated MCH74 and 7373 fibroblast cell lines *in vitro* with SSTC3 at 1μM for six or 24 hours (Figure 6). Qualitatively, bundles of MCH74 and 7373 treated with 1μM SSCT3 did not show any major differences (Figure 6A). In MCH74 fibroblasts, SSTC3 treatment reduced the level of vimentin bundles from 20.5±3.1% to 12.6±1.6% within three hours, with no further decrease past three hours. Conversely, treatment of 7373 fibroblasts with SSTC3 showed a progressive decrease in the percentage of the cells carrying IF vimentin bundles from a basal level of 46.5±2.2% to 27.6±1.3% at 24h (Figure 6B). This slight effect suggests that CK1α activation is insufficient to resolve the ARSACS phenotype.

**Figure 6.**
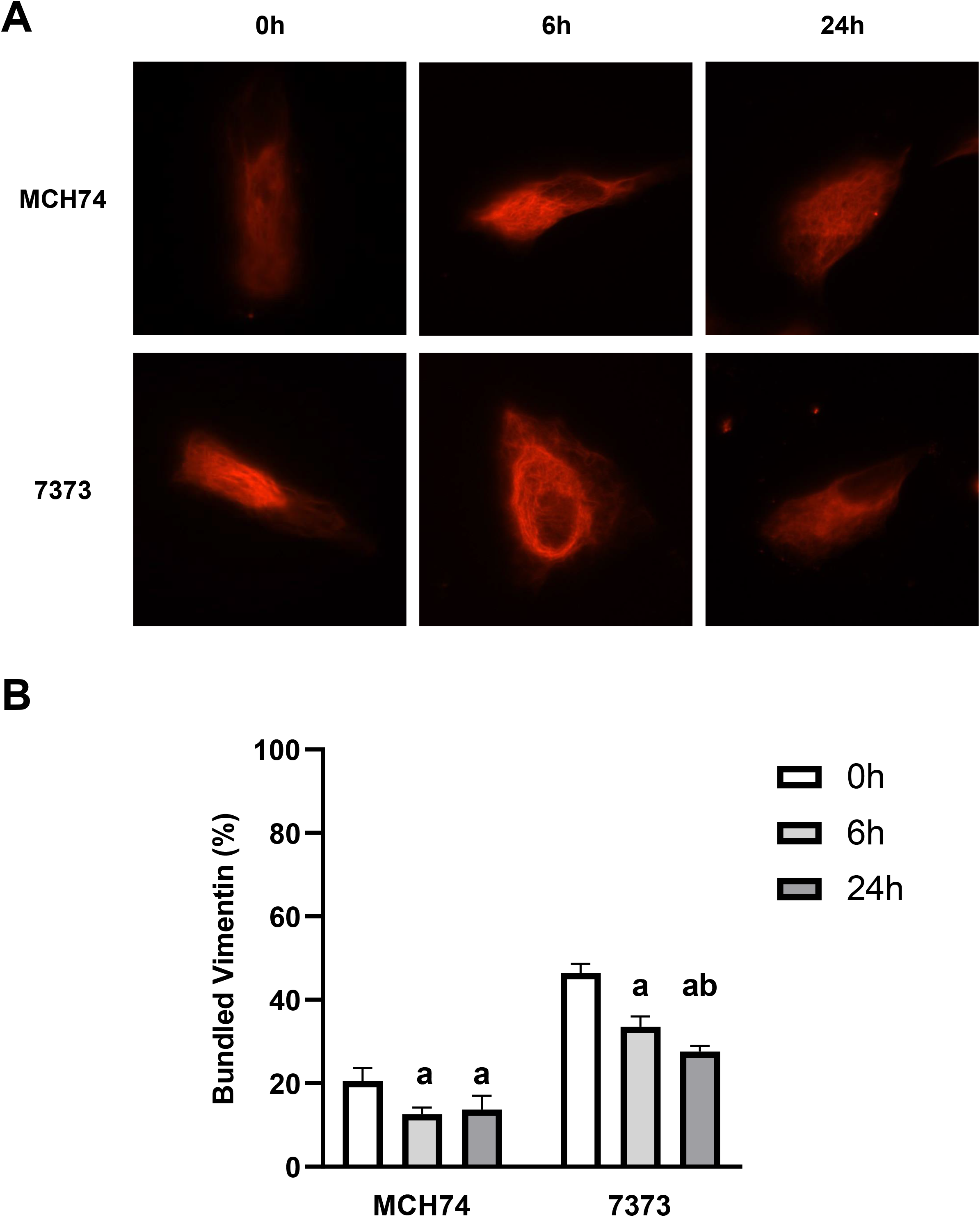
The effect of a CK1α activator SSTC3 on vimentin bundling in healthy and ARSACS human dermal fibroblasts. Human dermal fibroblast from control (MCH74) and ARSACS (7373) donors were treated with with 1 μM SSTC3, a CK1α activator, and the vimentin network was assessed via indirect immunofluorescence. **A**. Representative images of vimentin bundles in MCH74 and 7373 fibroblast lines being treated with SSTC3 at 0, 6, and 24 hours. **B**. Quantification of the effect of SSTC3 on vimentin bundling in fibroblasts. Shown are the percentage of cells presenting with normal vimentin network in the different experimental conditions. SSCT3 caused a partial reabsorption of vimentin bundles in both conditions. Data are expressed as means ± S.D (*n*=3 replicates/condition) with significant differences between timepoints within cell types labelled a = *p*<0.05 versus 0h and b = *p*<0.05 versus 6h, as determined by one-way ANOVA with Tukey’s HSD post-hoc analysis.

## Discussion

In our mouse models and ARSACS patient fibroblasts, FAM83H, FAM83B, and CK1α co-localize at the sites of IF bundles, suggesting that these proteins may play a role in ARSACS pathology. In particular, our co-localization studies suggest a cell-specific activity of CK1α, conferred by the relative expression of its FAM83 adaptor proteins. Previous studies have demonstrated that CK1α and FAM83H co-localize with IFs in colorectal and ameloblastoma cells (18,19), an interaction that was thought to be a mutually dependent process in IF organization. However, unlike CK1α, FAM83H is not expressed ubiquitously (31), as shown here with its low expression in Purkinje neurons, a key cell-type affected in ARSACS pathology (27,28). Specifically, FAM83B and CK1α co-localize with NF bundles in Purkinje neurons in the *Sacs*^-/-^ mice cerebellum, while FAM83B, FAM83H, and CK1α co-localize with vimentin IF bundles in fibroblasts derived from ARSACS patients. These data, taken together with the low levels of FAM83H in Purkinje neurons, show a cell-type specificity of the CK1α adaptors. Consistent with previous literature showing an effect of FAM83H on vimentin and keratins (18,19,32,33), FAM83H knockout induced the formation of NFL bundling in this study, demonstrating that NFs are also a target of FAM83H. However, the cell-specific expression of FAM83H (fibroblasts) and FAM83B (neurons), along with NFL bundles that formed upon FAM83B knockout, suggests that FAM83B is a specific adaptor of CK1α onto NF proteins in a neuronal context. Given that FAM83H, FAM83B, and CK1α levels were higher in cellular models of ARSACS, it is particularly striking to observe that these FAM83B and CK1α proteins strongly co-localize with NFL bundles in the Purkinje neuron dendrites. Along with the inappropriate targeting of NFL into the dendrites of Purkinje neurons in the molecular layer, rather than in the axons of the granular layer of the *Sacs*^-/-^ cerebellum, these findings suggest a sequestration of FAM83B and CK1α onto NFL bundles.

Alterations in NF organization have been well established in various ARSACS models (4,5,21). In the present study, we report that the NFL network is mislocalized in the *Sacs*^*-/-*^ cerebellum. Our findings demonstrate an attenuated expression of NFL in the *Sacs*^*-/-*^ cerebellar granular layer which mislocalizes to Purkinje neuron dendrites. This phenotype may reflect disrupted slow axonal transport of NFL along Purkinje neuron axonal projections (34). Consequently, the sparse NFL network surrounding Purkinje neuron axons may lessen the structural support that these axons receive, which could contribute to their degeneration (14). While slow axonal transport of NF is a complex process involving multiple interactors, phosphorylation has been shown to play a role (14). Ackerley et al. (2003) report that the phosphorylation of NFH inhibited its axonal transport (35), a finding that may be relevant to ARSACS pathology, given that NFH is hypophosphorylated in the *Sacs*^-/-^ brain and accumulates in Purkinje neuron dendrites (21).

In that context, CK1α is a key enzyme responsible for bundle formation and likely their resorption. CK1α induced the most bundling during *de novo* assembly of the NFL network in SW13^-/-^ cells, highlighting its key role in NFL bundle formation. Similarly, the inhibition of CK1α enzymatic activity by D4476 induced the formation of IF and NF bundles in MCH74 and *Sacs*^+/+^ motor neurons while exacerbating the NFL bundling in *Sacs*^*-/-*^ motor neurons. These results are consistent with the NFH dephosphorylation induced by CK1α inhibition in differentiated NB2a/d1 cells (15), similar to the phenotype observed in ARSACS (21). Despite the retained CK1α kinase activity in ARSACS, activation of CK1α by SSTC3 led to minimal improvement in the resorption of IF bundles in patient fibroblasts. All together these results suggest that CK1α, along with FAM83H or FAM83B, is recruited to NF bundles but its enzymatic activity cannot resolve these bundles. Although this kinase is constitutively active, CK1 phosphorylation is hierarchical and typically depends on the prior phosphorylation of its substrate by other kinases (36). In addition, the phosphorylation of certain CK1 substrates requires a “docking site”, a short sequence on the substrate that is distant from the phosphorylation site (37). Whether IFs contain docking motifs for CK1α, or if sacsin interacts with kinases upstream to CK1α, remains uncharacterized. Nevertheless, our findings in the present study refine our understanding of CK1α-mediated IF organization in ARSACS, highlighting a future research direction to determine how sacsin influences these multi-component pathways to CK1α phosphorylation. Taken together, our data suggest that a priming event upstream of CK1α activation is compromised in ARSACS, impeding the ability of CK1α to remedy NF bundling.

In summary, our findings suggest that a priming event upstream of CK1α activation is compromised in ARSACS, impeding the ability of CK1α to remedy NF bundling. We propose that despite the appropriate recruitment of CK1α to NF bundles, a disrupted FAM83B protein sequesters CK1α at these sites. Therefore, these NF bundles would function as a ‘sink’ for FAM83H/B and CK1α proteins, contributing to the ARSACS molecular cascade. While FAM83B overexpression has been identified as an oncogene that facilitates cell transformation, migratory and invasive capabilities through the activation of various signaling pathways (20,38), why FAM83B is expressed in post-mitotic cells like Purkinje neurons remains unknown. Thus, understanding the relation of FAM83B in ataxia presents novel challenges.

## Acknowledgements

We thank Lai GT and Billette B for technical support.

## CRediT Author Contributions Statement

**Caitlin Atkinson:** Formal Analysis; Investigation; Visualization; Writing – original draft; Writing – review & editing; **Zacharie Cheng-Boivin:** Formal Analysis; Investigation; Visualization; Writing – original draft; Writing – review & editing; **Nancy Larochelle:** Supervision; **Sandra Minotti:** Resources; **Benoit Gentil:** Conceptualization; Funding Acquisition; Methodology; Project Administration; Supervision; Visualization; Resources; Writing – original draft; Writing – review & editing.

