## Supplemental Material for "CK1α, FAM83H, and FAM83B contribute to bundling of neurofilaments and are sequestered in cellular and mice models of ARSACS"

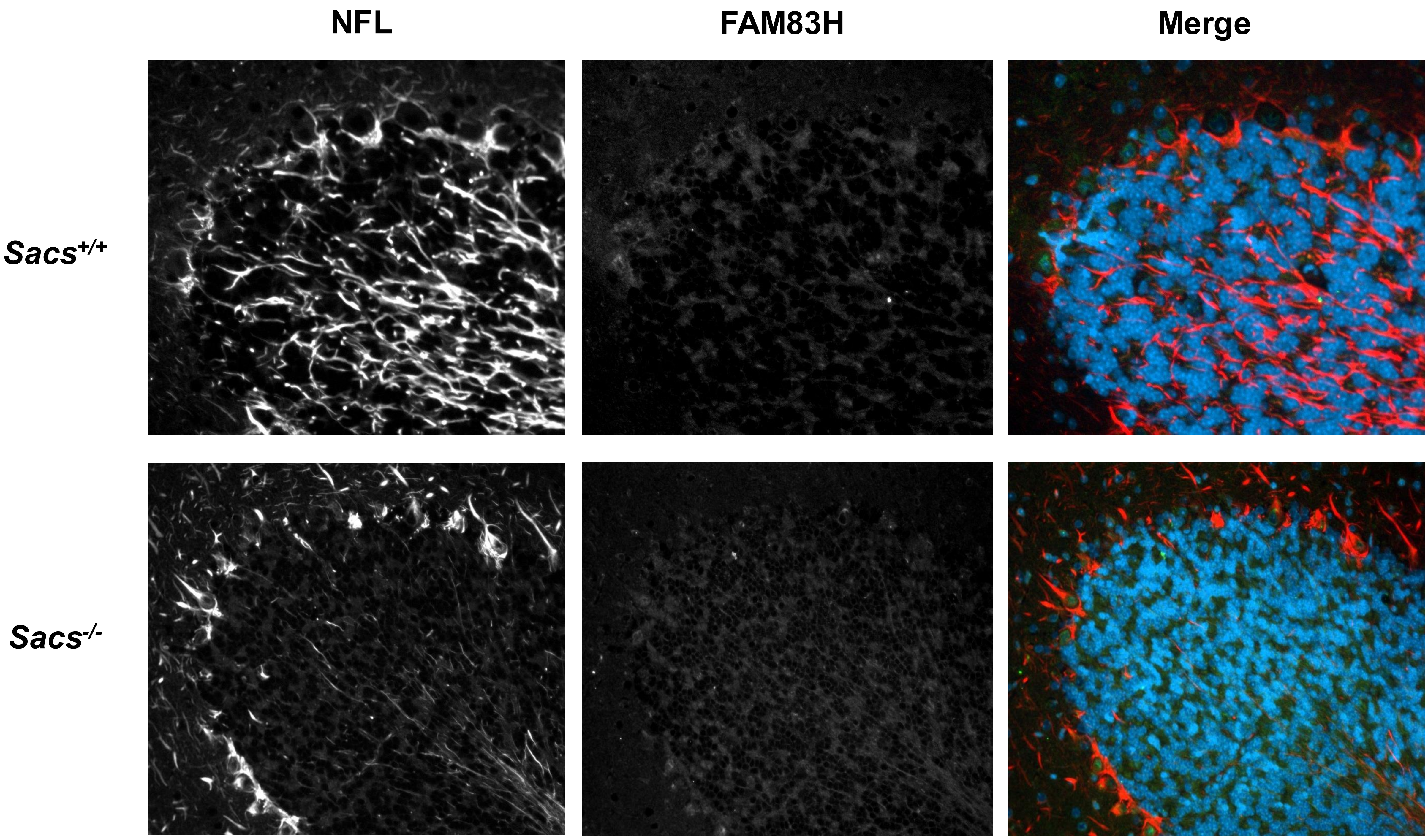


Supplementary Data 1. Expression FAM83H in the Sacs+/+ and Sacs-/- adult mouse cerebellum. FAM83H was not detected in the cerebellum.

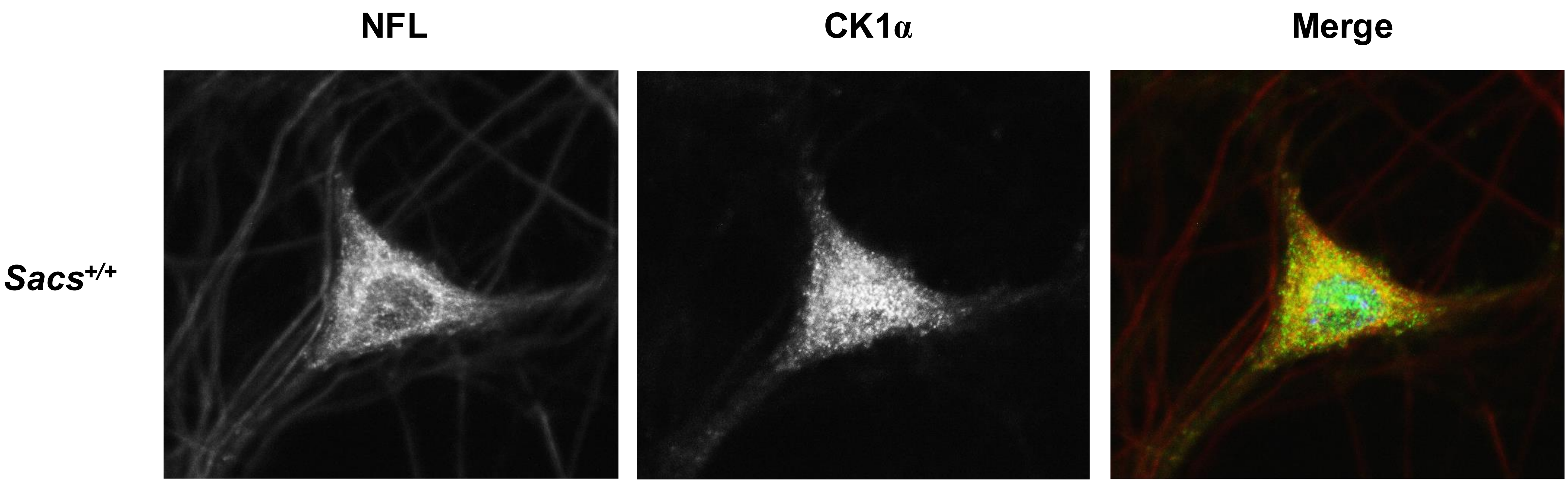


Supplementary Data 2. The expression of CK1α in cultured motor neurons. CK1α was expressed and co-localized with NFL in motor neurons.

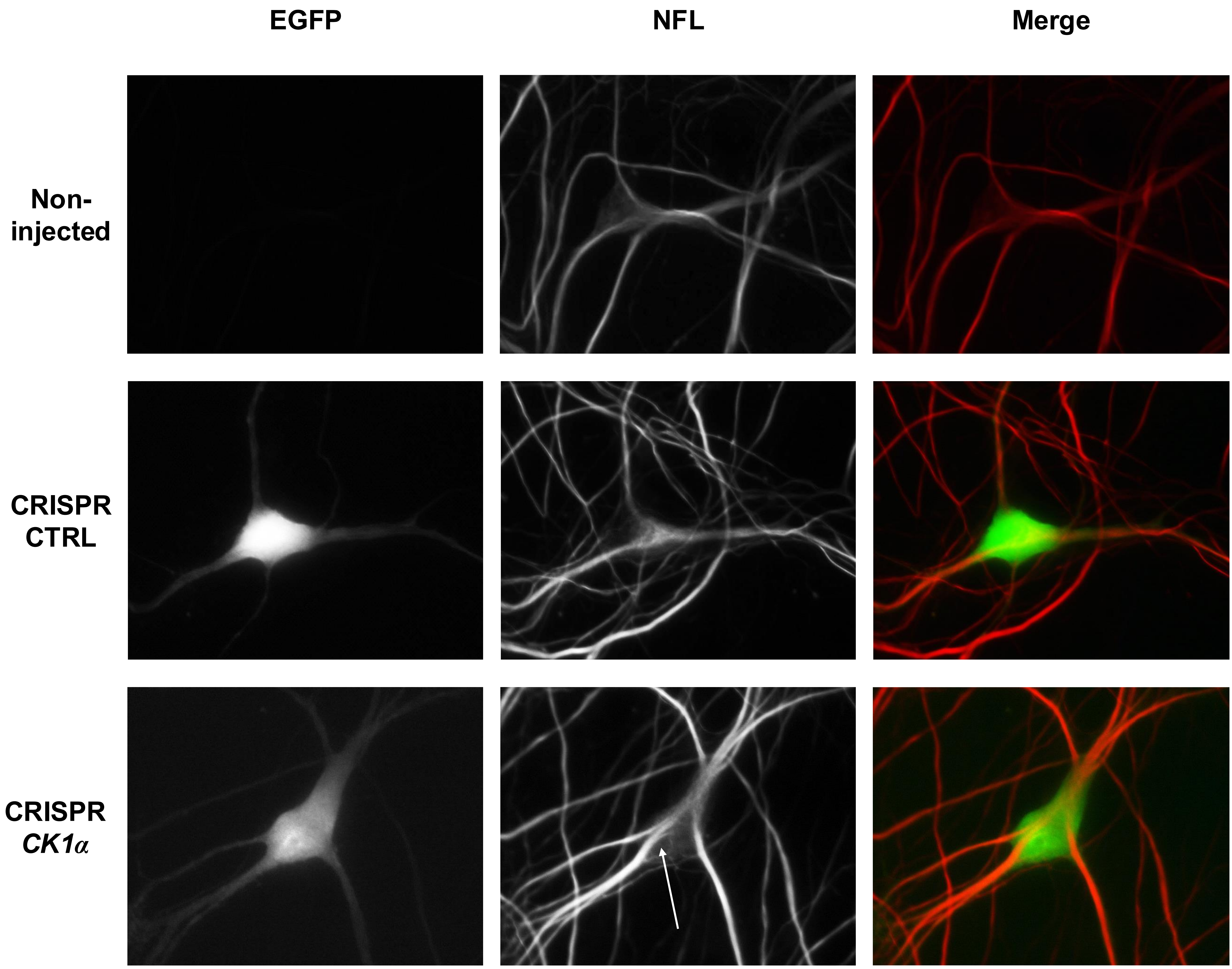


Supplementary Data 3. The effect of CK1α knockout in cultured motor neurons. Motor neurons in spinal cord-dorsal root ganglia cultures were micro-injected with a CRISPR-Cas9 knockout plasmid carrying a control guide RNA or guide RNA targeting CK1α and expressing EGFP. Images show immunolabelling for NFL network. The arrow highlights the abnormal bundling of NFL with CRISPR-CK1α, resembling that of a Sacs-/- cells, which is absent in non-injected and CRISPR-CTRL neurons.
